# Delving into the claustrum: insights into memory formation, stabilization and updating in mice

**DOI:** 10.1101/2024.04.26.591314

**Authors:** Candela Medina, Santiago Ojea Ramos, Amaicha M Depino, Arturo G Romano, María C Krawczyk, Mariano M Boccia

## Abstract

The claustrum is a brain structure that remains shrouded in mystery due to the limited understanding of its cellular structure, neural pathways, functionality and physiological aspects. Significant research has unveiled connections spanning from the claustrum to the entire cortex as well as subcortical areas. This widespread connectivity has led to speculations of its role in integrating information from different brain regions, possibly contributing to processes such as attention, consciousness, learning and memory. Our working hypothesis posits that claustrum neural activity contributes to the formation, stabilization and updating of long-term memories in mice. We found evidence in CF-1 mice of a decline in behavioral performance in an inhibitory avoidance task due to intra-claustral administration of 2% lidocaine immediately after a training session or memory recall. Nevertheless, this does not seem to be the case for the acquisition or retrieval of this type of memory, although its neural activity is significantly increased after training, evaluated through c-Fos expression. Moreover, inhibition of the claustrum’s synaptic activity appears to impair stabilization but not the acquisition or retrieval of an unconditioned memory formed in a nose-poke habituation task.

## 1. Introduction

Memory, a fundamental cognitive process, captivates researchers as they seek to understand its mechanisms underlying formation, consolidation and retrieval ^1^. Within the complex network of the human brain, the claustrum—a nucleus nestled deep within the cerebral cortex—has emerged as a subject of profound interest for neuroscientists studying memory and its neuropharmacology ^2,3^. This research paper aims to study the participation of the claustrum in memory processes in mice.

The claustrum (CLA), or “hidden place” in Latin, is a thin, irregular sheet of gray matter bilaterally positioned along the insular cortex ^4^. Structurally, it assumes an elongated shape that spans the rostrocaudal axis of the brain, extending beneath the neocortical mantle and establishing connections with the putamen, globus pallidus and the extreme capsule ^5^. This positioning emphasizes its potential role as an integrative hub within the brain’s vast neuronal network ^6^.

The CLA’s connectivity further highlights its significance, as it possesses extensive reciprocal connections with diverse cortical and subcortical regions. This connectivity allows it to form a network that extends throughout the entire cerebrum ^7^. Noteworthy, these connections encompass the prefrontal cortex, motor cortex, somatosensory cortex, as well as the visual and auditory association cortices ^8^. Additionally, subcortical connections extend to the basal ganglia ^5^, thalamus ^9,10^, amygdala ^5,11^ and hippocampus ^12,13^, among other regions ^5^. Such extensive connectivity underscores the CLA’s capacity for coordinating information flow across diverse brain regions, suggesting a role in integrating complex sensory, motor and cognitive processes ^3,14,15^.

Despite decades of investigation, the precise functions of the CLA remain elusive. Its structural characteristics, extensive connections and neurochemical profile position it as a prime candidate for involvement in attention, consciousness, perception and decision-making processes ^3,16^. Moreover, emerging evidence suggests its contribution to memory formation and retrieval ^17–21^, together with interactions with memory-related regions, such as the hippocampus and prefrontal cortex ^21,22^. Additionally, its dynamic connections with the limbic system and its role in mediating attention and sensory integration further support its involvement in memory function ^16^.

The abundance of functions that the CLA appears to exert is associated with a wide variety of potential dysfunctions. Due to the preponderant role it seems to play in higher cognitive processes, researchers are investigating its involvement in various neurological pathologies, including Wilson’s disease ^23^, epilepsy ^24^, autism ^25^, depression, schizophrenia ^26^ and Lewy body dementia ^27^, as well as neurodegenerative diseases such as Alzheimer’s ^28^ and Parkinson’s ^29^.

Memory consolidation and reconsolidation processes are essential for the stabilization and modification of memories ^30–33^. Consolidation entails the transformation of newly acquired information into long-term memory through cellular and molecular events such as synaptic plasticity, gene expression and protein synthesis ^34^. Similarly, memory reconsolidation occurs when a previously consolidated memory is retrieved and subsequently re-stabilized, allowing for potential updates or modifications ^35–37^. Given the CLA’s extensive connections and its involvement in memory-related processes, it emerges as a compelling candidate for influencing both consolidation and reconsolidation phenomena.

In the current investigation, our primary goal is to explore the involvement of the CLA in memory consolidation and reconsolidation processes in mice. To accomplish this, we employed two different behavioral tasks, each presenting diverse motivational and motor challenges. By employing these tasks, we aim to shed light on the CLA’s crucial role in learning and memory.

## 2. Results

### 2.1 The claustrum’s role in the acquisition and consolidation of an associative memory

We investigated claustral neural activity following inhibitory avoidance (IA) conditioning by assessing c-Fos expression via immunohistochemistry (IHC; ^44^) after the training trial (median step-through latency = 14 sec, IQR = 12.75 - 18.25 sec). Behaviorally naïve animals served as a control group (Figure 1A). We also included a cohort of behaviorally naïve animals, which served as a control group for the trained set of mice (Figure 1A). To delineate the boundaries of this brain structure, we relied on the concentrated clustering of parvalbumin (PV) labeling, which serves as a defining feature of the CLA ^45,46^. On top of claustral neural activity, we evaluated c-Fos expression in the CA1 subarea of the dorsal hippocampus. Numerous studies have suggested that hippocampal subregions are involved in different mechanisms within mnemonic processing (reviewed in ^47^), including evidence of impairments observed in contextual memory processing that not only appear to depend specifically on the dorsal hippocampus, but more particularly on the CA1 subregion ^48^. Furthermore, we investigated neural activity in the motor cortices (M1/2) of the mice, brain regions not previously associated with the processing and encoding of associative memories (Figure 1A and B).

**Figure 1.**
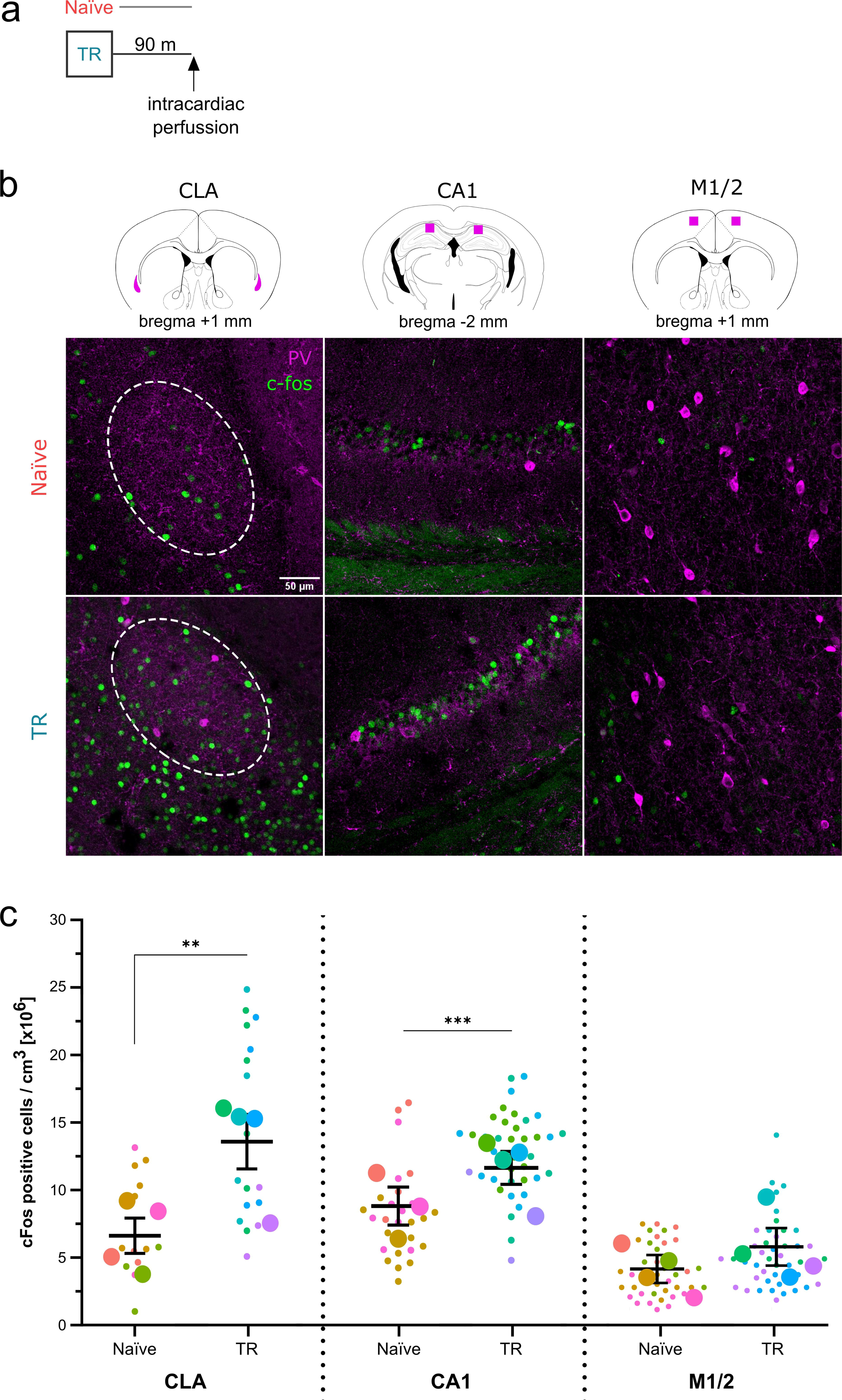
Neuronal activity induced by IA training. **(A)** We performed IHC on brain sections 90 min after IA training to reveal c-Fos expression in the claustrum (CLA, droplet-like areas in bregma +1 mm slice), CA1 subarea of the dorsal hippocampus (squared areas in bregma −2 mm slice) and motor cortices (M1/2, squared areas in bregma +1 mm slice). We also performed IHC for PV in order to delineate the claustral area in each brain slice. **(B)** Representative CLA, CA1 and M1/2 sections from brain slices of trained (TR) and Naïve mice. The dotted line represents the area evaluated as CLA according to PV IHC. Scale bar represents 50 μm. **(C)** c-Fos density for Naïve and TR animals, evaluated in CLA, CA1 and M1/2 brain slices (n_Naïve_ = 2-13 slices from 4 mice, n_TR_ = 2-14 slices from 4 mice). Small dots represent each brain slice, big dots represent each animal’s mean c-Fos density and the group’s mean and standard deviation are represented in black lines.

As shown in Figures 1B and C, both claustral (CLA, left panel) and hippocampal (CA1, central panel) neural activity significantly increased after the IA training trial (p < .01 and .001, respectively; exp. 1 in Suppl. Table 1). Conversely, the learning experience did not significantly alter c-Fos expression in the motor cortices (M1/2, right panel; p > .05; exp. 1 in Suppl. Table 1).

Given the significant upregulation of claustral neural activity following the IA training trial, we next investigated its role in associative memory acquisition. We administered bilateral intra-claustral infusions of either vehicle (Veh) or 2% lidocaine (Lido) solution 1 hour prior to the IA training trial to inhibit synaptic transmission in the claustrum during memory formation (Figure 2A). Lidocaine blocks voltage-gated sodium channels, leading to a reversible block of action potential propagation ^49^. As shown in Figure 2B, claustral synaptic inhibition before training did not significantly affect the behavioral performance of Lido-treated animals compared to control mice (p > .05, exp. 2 in Suppl. Table 1).

**Figure 2.**
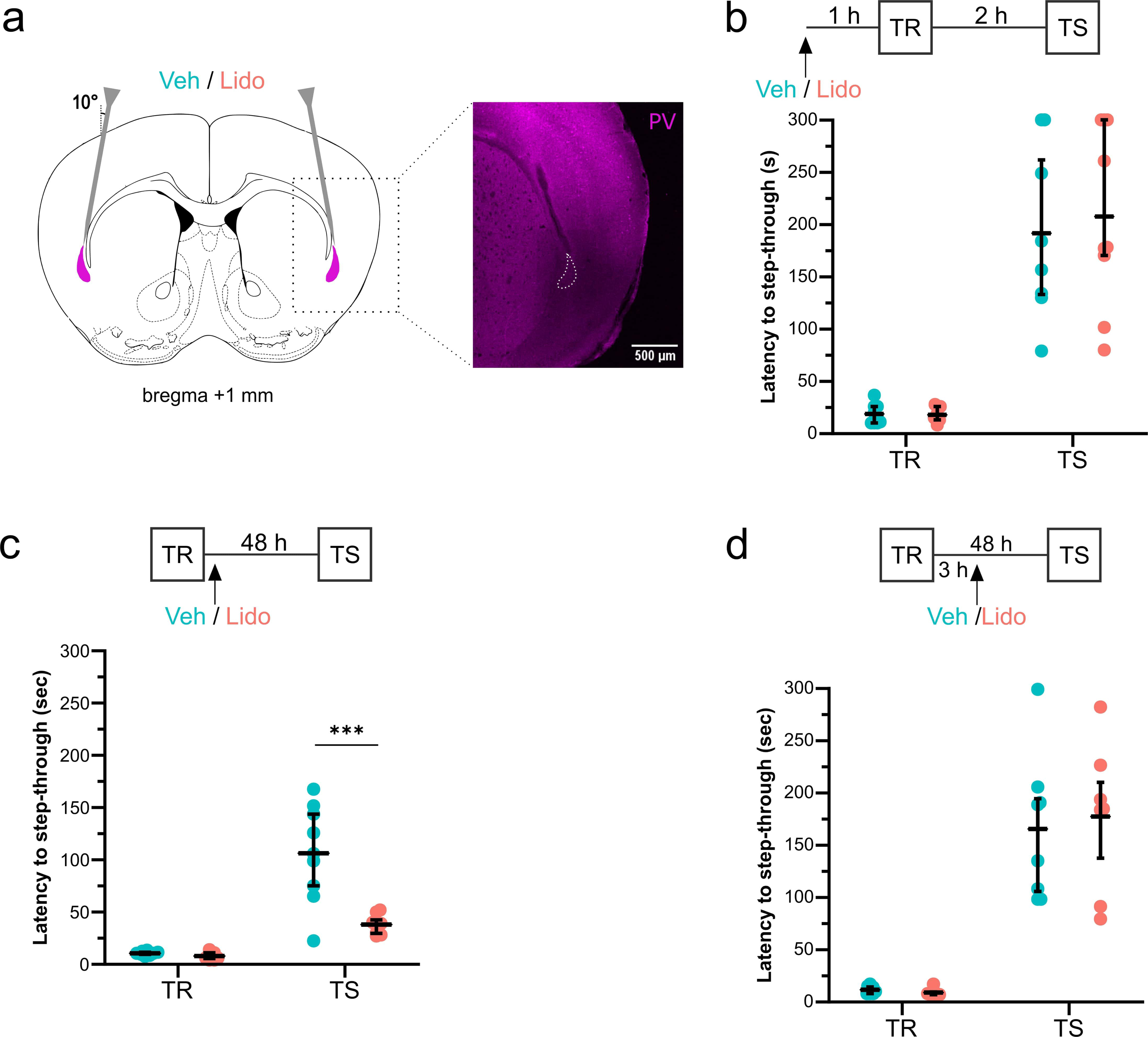
The claustrum in the acquisition and consolidation of an associative memory. **(A)** We bilaterally infused the CLA with either a 2% lidocaine (Lido) or saline solution (Veh) into each hemisphere. Inset: IHC for PV, showing the needle trajectory. Scale bar represents 500 μm. **(B-D, top panel)** We injected mice in the CLA with Veh or Lido solution either **(B)** 1 h prior to the training trial (TR; n_Veh_ = 8, n_Lido_ = 9), **(C)** immediately after TR (n_Veh_ = 9, n_Lido_ = 8) or **(D)** 3 h after TR (n_Veh_ = 8, n_Lido_ = 7). The retention test took place either 2 or 48 h later (TS). **(B-D, bottom panel)** Step-through latencies of animals during TR and TS trials. Median values and IQR are shown in black, individual values in color.

As previously mentioned, memory consolidation entails the stabilization of newly acquired information into long-term memory ^50^. To address the role of the CLA in this memory process, we trained two groups of mice and, immediately after training, infused their CLA with either a Veh or Lido solution. After 48 hours, we assessed their behavioral performance (Figure 2C). After 48 h, their behavioral performance in the task was assessed, as shown in Figure 2C. In contrast to control mice administered with Veh, which presented a higher latency to step-through to the dark compartment, inhibition of claustral synaptic transmission by Lido significantly impaired the performance of mice in this task (p < .0001; exp. 3 in Suppl. Table 1).

In order to evaluate whether this impairing effect of Lido on long-term memory was time-specific, we repeated the experiment with another cohort of mice, except for administering either Veh or Lido solutions 3 h after the training trial. As shown in Figure 2D, delayed infusions did not affect the step-through latencies in either group (p > .05, exp. 4 in Suppl. Table 1). These last experiments, depicted in Figures 2C and D, were replicated in female mice, yielding similar results (Suppl. Figure 1A and B, respectively).

Considering the results of this first set of experiments, it appears that the inhibition of claustral synaptic transmission impairs the consolidation of associative memory without significantly affecting its acquisition, despite a notable increase in neural activity after training.

### 2.2 The CLA in the acquisition and consolidation of an habituation memory

Next, we wanted to evaluate whether the CLA is relevant for the acquisition and stabilization of another type of memory, entailing different motivational, sensorial and motor requirements. We trained two groups of mice in a nose-poke habituation task (HT) by exposing them to a hole-board for 5 minutes, recording cumulative nose pokes over time (min 1: p = .7791, min 2: p = .6982, min 3: p = .5863, min 4: p = .1679, min 5: p = .0540; exp. 5 in Suppl. Table 1; left panel of Figure 3A). Immediately after this training trial, we administered bilateral infusions of either Veh or Lido into the CLA of two groups of mice. During the testing trial, mice treated with Lido exhibited reduced cumulative nose pokes compared to the control group, indicating impaired habituation to the stimulus (min 1: p = .3962, min 2: p = .3274, min 3: p = .0407, min 4: p = .0550, min 5: p < .0010; exp. 5 in Suppl. Table 1; right panel of Figure 3A).

**Figure 3.**
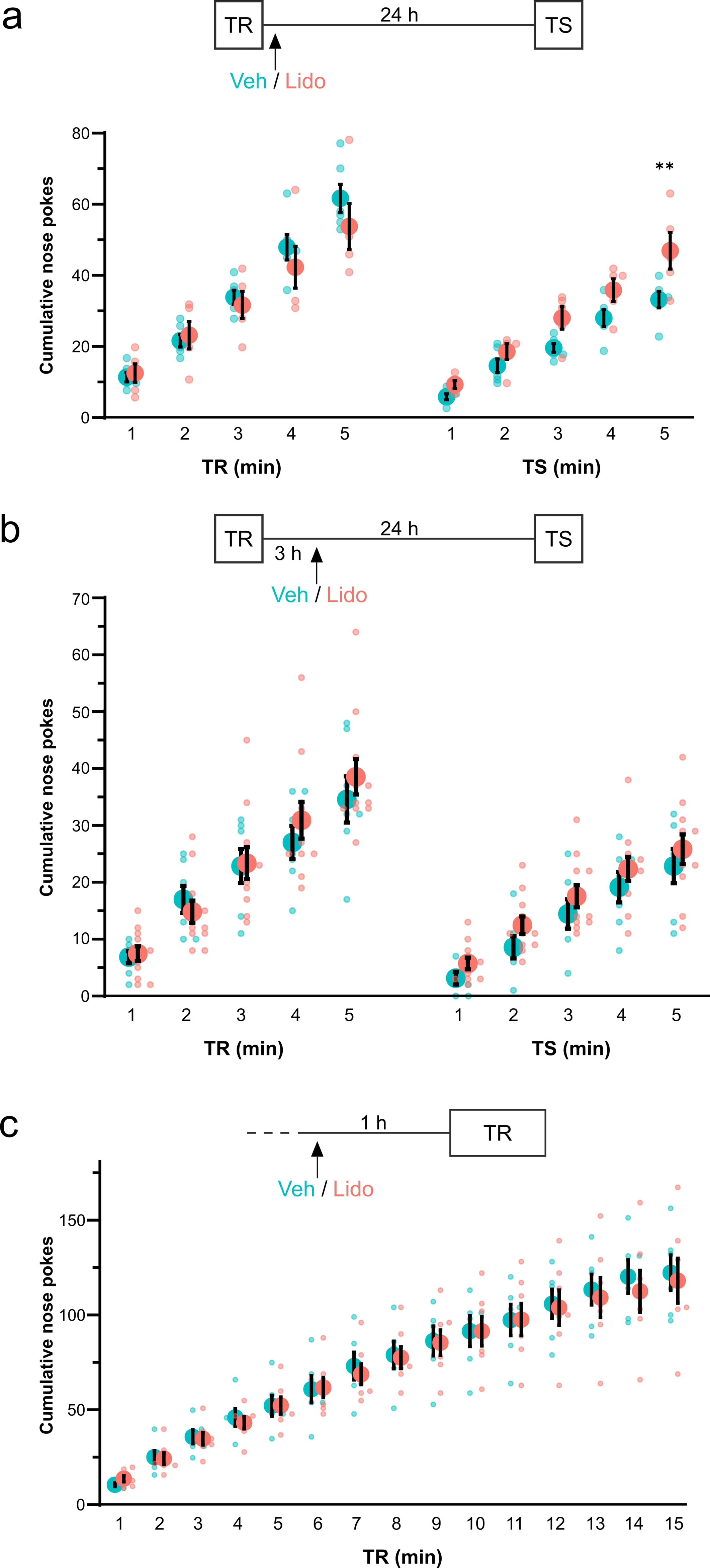
The claustrum in the acquisition and consolidation of an habituation memory. **(A-C, top panel)** We injected mice in the CLA with Veh or Lido solution either **(A)** immediately to the training trial (TR; n_Veh_ = 6, n_Lido_ = 5), **(B)** 3 h after TR (n_Veh_ = 7, n_Lido_ = 11) or **(C)** 1 h prior TR (n_Veh_ = 7, n_Lido_ = 11) of a nose-poke habituation task. When indicated, the retention test took place 24 h later (TS). **(A-C, bottom panel)** Cumulative nose pokes performed by animals through time during TR and TS trials. Big dots represent the group’s mean values, small dots stand for individual values and standard errors are shown in black.

Next, we tested whether the effect of intra-claustral Lido administration on long-term habituation was time-specific. When synaptic transmission in the CLA was impaired 3 hours after training, the animals’ performance in the task did not significantly differ from the control mice (p > .05, exp. 6 in Suppl. Table 1; Figure 3B).

As before, we also investigated whether claustral synaptic activity was necessary for the acquisition of this type of memory by administering either Lido or Veh 1 hour before the training trial, extending the trial duration to 15 minutes to reveal possible intra-trial habituation. Again, there were no significant differences in task performance between Lido- and Veh-administered mice (p > .05, exp. 7 in Suppl. Table 1; Figure 3C).

Altogether, this last set of results point to an essential role of claustral synaptic activity in long-term storage of an habituation memory, whereas this area does not seem to be required for the acquisition of this type of memory.

### 2.3 The CLA in the retrieval and labilization/reconsolidation of an associative memory

Next, we aimed to address the role of the CLA in memory retrieval. We administered either a Veh or Lido solution 1 hour prior to re-exposure to the illuminated platform to prevent synaptic transmission in this area during recall of the associative memory (TS1 in Figure 4A). When tested 24 h later, we observed that claustral synaptic inhibition does not cause a significant effect on the behavioral performance of Lido-administered animals, when compared to control mice (p > .05, exp. 8 in Suppl. Table 1; Figure 4A).

**Figure 4.**
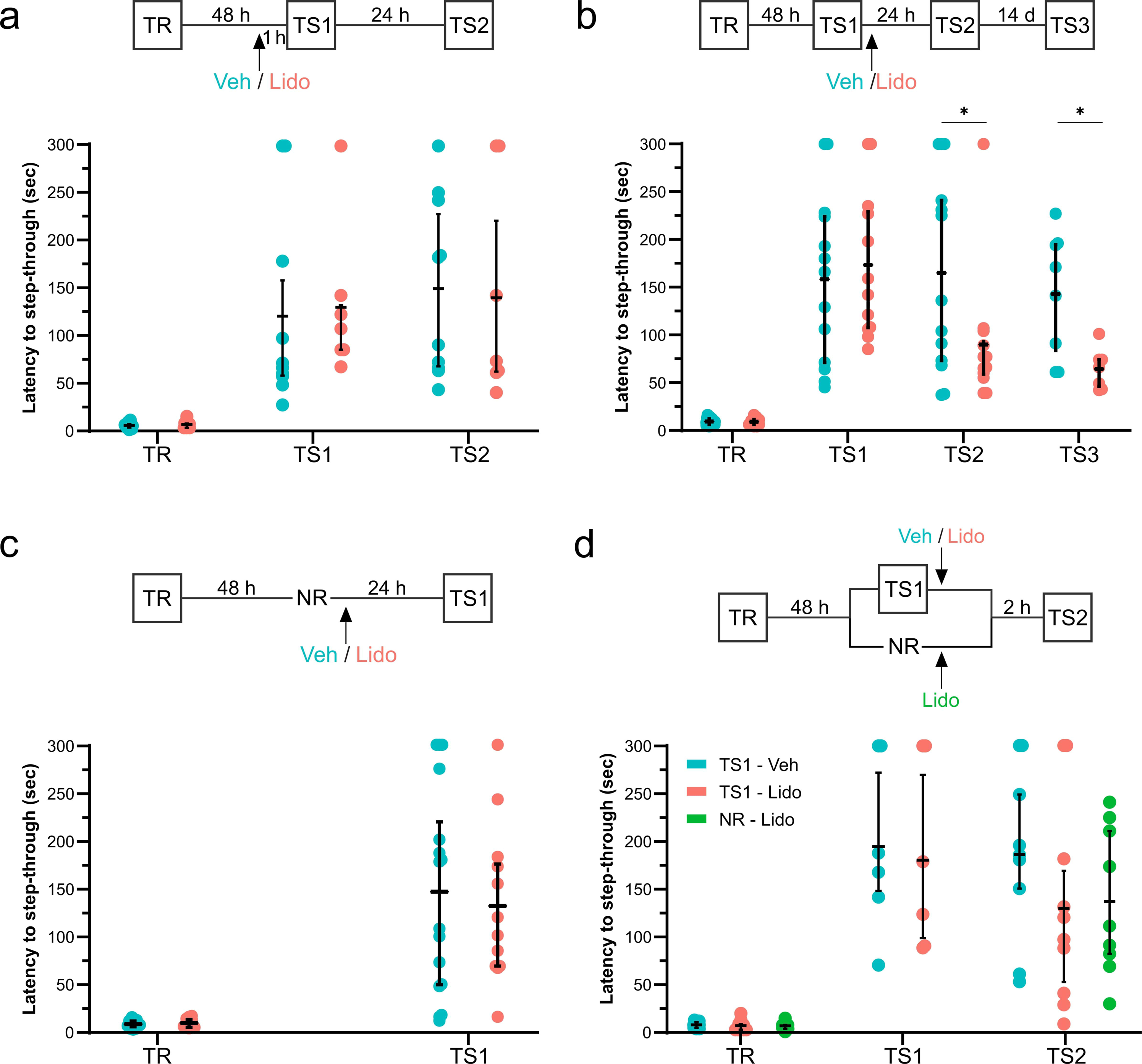
The claustrum in the labilization/reconsolidation of an associative memory. **(A-D, top panel)** We injected mice in the CLA with Veh or Lido solution either **(A)** 1 h prior to the first retention test (TS1; n_Veh_ = 10, n_Lido_ = 7), **(B)** immediately after TS1 (n_Veh_ = 13, n_Lido_ = 12), **(C)** 48 h after TS1 (n_Veh_ = 8, n_Lido_ = 6) or **(D)** immediately after TS1 (n_TS1-Veh_ = 9, n_TS1-Lido_ = 10) and the third was administered with Lido 48 h after training (n_NR-Lido_ = 9). When indicated, the second retention test took place either 2 or 24 h later (TS2) and a third test was performed after 14 d (TS3). **(A-D, bottom panel)** Step-through latencies of animals during training, TS1, TS2 and TS3 trials. Median values and IQR are shown in black, individual values in color.

Following a reactivation-eliciting reminder, a memory trace can undergo the processes of labilization and reconsolidation, ultimately leading to its re-stabilization with potential updates or modifications ^35–37^. We aimed to study the CLA’s role in these processes using the IA task. We trained two groups of mice and evaluated their behavioral performance in the task 48 h later by exposure to the illuminated platform, a reminder that has proven to entail memory reactivation (TS1 in Figure 4B; ^51^). Immediately after TS1, we performed bilateral intra-claustral infusion of either a Veh or Lido solution and re-tested their performance 24 h later (TS2). The inhibition of claustral synaptic transmission with Lido significantly hindered the mice’s performance in this task, when compared with the control mice given Veh (p = .0141, exp. 9 in Suppl. Table 1) and this effect was still present up to 14 d later (p = .0128, exp. 9 in Suppl. Table 1; TS3 in Figure 4B). On top of that, this impairment is specific to recall, since Lido-injected mice did not differ in their IA performance to the control animals, when infused 48 h after training and tested 24 h later (p > .05, exp. 10 in Suppl. Table 1; Figure 4C).

As previously mentioned, memories are dynamic and subject to influence by various processes like reconsolidation and extinction, which can alter their strength or content ^52,53^. These processes operate through distinct mechanisms, resulting in different outcomes. Reconsolidation allows for the integration of new information and potential strengthening of the memory, while extinction leads to the formation of a new memory trace with a behavioral response opposite to the original one ^54^. In this case, the formation of an extinction memory would result in a lower latency to step-through to the dark compartment. Therefore, we aimed to determine whether the diminished behavioral performance observed when synaptic transmission was inhibited after recall (Figure 4B) was either due to an impairment in the reconsolidation of the original memory or a facilitation in the formation of a new extinction memory. Similarly as before, we administered two groups of animals with either Veh or Lido in the CLA immediately after TS1, but evaluated their performance in the task 2 h later, to disclose any possible short-term differences in their behavior. Lido-infused mice did not present significant differences in their step-through latencies compared to control animals, nor did drug-administered mice which did not undergo memory recall (p > .05, exp. 11 in Suppl. Table 1; NR-Lido in Figure 4D).

Altogether, the results of this last set of experiments point to a destabilizing effect on memory of claustral inhibition after its retrieval, probably due to an impairment in the reconsolidation process. However, the CLA does not seem to have a crucial role in the retrieval of this associative memory.

## 3. Discussion

In the present study, we provide new evidence for the central role of the claustrum in memory processes, particularly consolidation and reconsolidation. Through behavioral tasks and pharmacological interventions, we demonstrate a correlation between claustral neural activity and memory-related processes, showing that this activity is crucial for memory storage and updating.

First, both the CLA and the CA1 area of the hippocampus showed an upregulation of the expression of the neural activity marker, c-Fos, after training. This suggests that the CLA, as it has been proven for CA1 ^55^, takes part in the information processing related to the learning experience. It has been previously documented that c-Fos expression is triggered in numerous brain regions after either a learning event or memory reactivation ^44^ and, of significance, that exposure to a novel environment has been shown to induce c-Fos expression in a significant portion of claustral neurons ^56–58^. On top of that, CLA activity has been correlated to the acquisition of a delayed eye-blink response ^18^ and cocaine conditioned-place preference ^20^.

Since immunohistochemistry fails to reveal whether the increase in c-Fos expression is a result from learning, memory consolidation or both, we opted for a pharmacological approach. Intracerebral infusion of lidocaine, known for its ability to prevent propagated action potentials through reversible blockade of voltage-gated Na+ channels ^59^, has been extensively used as an effective method for examining the functional involvement of neural structures in behavior ^60–62^. Performance impairment was observed only when lidocaine was administered immediately after IA training, suggesting the CLA’s critical role in memory storage for this task. To further explore memory processes, we used a nose-poke habituation task (HT) that requires different motivational, motor, and emotional states compared to the IA task. Performance impairment was again present only when lidocaine was administered immediately after HT training, suggesting a certain degree of generalization in the types of memories that rely on claustral activity for this process.

It is worth mentioning that the CLA is crossed by fibers originating from the frontal, occipital, and temporal lobes ^63^. Thus, inhibiting its neural activity with lidocaine could complicate result interpretation. However, our findings align with those of Terem and colleagues, indicating that the activity of claustral neurons expressing dopamine D1 receptors is essential for establishing cocaine conditioned-place preference and can induce place preference via optogenetic stimulation ^20^. Additionally, recent evidence supports that CLA stimulation during non-REM sleep enhances labile associative memory consolidation ^17^.

Strikingly, under our experimental conditions, local lidocaine infusion before training did not affect IA performance nor HT acquisition rates, suggesting a transient effect not impacting attentional or acquisition processes. This contrasts with findings linking CLA activity to delayed eye-blink response acquisition ^18^ and the role of claustral neurons expressing dopamine receptors in acquiring cocaine conditioned-place preference ^20^. Recent research indicates that the claustrum enhances distraction resilience and modulates vigilance in response to anxiety-related threats ^64,65^. Anatomical studies show robust inputs from limbic structures, like the basolateral amygdala ^64,66^, conveying salience signals, positioning the CLA as a limbic-sensorimotor interface ^65^. However, our findings contradict this: CLA inactivation during training did not impair performance, suggesting salience processing is independent of claustral activity in our conditions. Nevertheless, without measuring the local concentration of lidocaine in the CLA 60 minutes after administration, its potential impact on neural activity remains uncertain. Studies in monkeys showed neuronal shutdown occurring within 5 minutes of starting the injection and recovering within 30 minutes ^67^. However, considering its reported pharmacokinetics and half-life (1.2 ± 0.3 hours after intravenous administration in rats ^68^), together with the local administration route, we can infer that lidocaine was likely still present during training. Further studies are required to confirm this assumption.

When a subject is exposed to a reactivation session, it triggers at least one of two memory processes: reconsolidation and/or extinction ^69^. The first allows updating and/or strengthening of the trace ^35–37^, while the last leads to the formation of a new memory with a behavioral response opposite to the original one ^54^. In the case of the IA task, extinction would result in the animal entering the dark compartment more quickly, leading to a shorter latency to step-through. In our experiments, latencies to step-through decreased only when lidocaine was administered immediately after reactivation, without affecting retention latencies in the absence of a reactivation session or long-term memory consolidation processes. Using a spontaneous recovery protocol ^69^, we found no memory recovery up to 17 days post-acquisition. Additionally, treated mice tested 2 hours after reactivation showed no extinction memory formation. These results suggest the claustrum’s critical role in the memory reconsolidation of an IA task, providing initial evidence of its specific involvement in memory reactivation and restabilization. Conversely, claustral activity may not be essential for memory retrieval, as mice performance remained unaffected when the pharmacological treatment was administered before reactivation. This aligns with findings by Terem and colleagues, who reported that claustral neuron activity is crucial for the acquisition of cocaine conditioned-place preference but not for its expression ^20^.

To sum up, our study provided novel evidence suggesting that the claustrum, an often overlooked brain structure, is crucial for the storage and updating of memories. Additionally, our findings reveal a correlation between claustral neural activity and memory-related processes, as evidenced by the upregulation of c-Fos expression following training in an IA task. These findings may hold translational significance for neurological pathologies and neurodegenerative diseases, potentially offering new insights for therapeutic interventions to mitigate cognitive decline. This research deepens our understanding of memory mechanisms and underscores the claustrum’s importance in cognitive functions.

## 4. Methods

### 4.1 Animals

We used 223 CF-1 mice from our own breeding stock (age: 60–70 d; weight: 25–30 g; 190 male and 33 female). They were housed in groups of 5-10 mice per cage (cage size: 50 × 30 × 15 cm) with food and water freely available, and were kept in a 12:12-hour light–dark cycle (lights on at 06.00 h) in a temperature regulated (23–25 1C) environment. We conducted the experiments between 08.00 and 14.00 h and carried them out in accordance with the National Institute of Health Guide for the Care and Use of Laboratory Animals (NIH Publication N1 80– 23/96) and local guidelines establish by CICUAL-FFyB (CUDAP # 0044975-2016). All efforts were made to minimize animal suffering and to reduce the number of animals used. The number of animals used is stated for each experiment in Supplementary Table 1.

### 4.2 Stereotaxic surgery and intraclaustral injections

We anesthetized mice with isoflurane and secured them in a stereotaxic apparatus. The surgical procedure involved making an incision in the scalp and, to access the claustrum (CLA), we drilled a hole in the skull of each hemisphere at the following stereotaxic coordinates: AP +1.0 mm relative to bregma, L ±2.8 mm from the midsagittal suture. Then, we lowered a 30 G stainless-steel needle into each hemisphere in a 10 degree angle from the sagittal plane (−3.8 mm from the skull surface, Figure 1A; ^38^) and bilaterally infused the claustrum with either a 2% lidocaine or saline solution (see Drugs section). Injections were driven by an automatic pump (MD-1020, Bioanalytical Systems, Inc.) at a rate of 0.52 uL/min through the needle attached to a 5 μl Hamilton syringe with PE-10 tubing. The volume of each infusion was 0.2 μl/hemisphere. We closed the incision with a stitch and the animals were returned to their home cages.

We determined the accuracy of the injections by histological examination of the needle position on an animal-by-animal basis. For this purpose, we quickly euthanized animals by cervical dislocation followed by decapitation. Next, we dissected and fixed their brains in 4% paraformaldehyde in buffer phosphate saline. Then, we cut them into 200 nm coronal sections with a vibratome. The deepest position of the needle was superimposed on serial coronal maps (Franklin & Paxinos, 1997). Animals were excluded from the statistical analysis if the infusions had caused excessive damage to the cerebral tissue or if the needle trajectory extended outside of the claustrum.

### 4.3 Drugs

We bilaterally administered a solution of 2% lidocaine (Lido; Laboratorios Veinfar I.C.S.A.. Bs. As) or saline (Veh). We determined this Lido dose based on previous work from McIntyre and collaborators ^39^. All other agents were of analytical grade and obtained from local commercial sources.

### 4.4 Behavioral Procedures

#### 4.4.1 Inhibitory Avoidance task

We studied inhibitory avoidance (IA) behavior in a one-trial learning, step-through type situation ^40,41^, which utilizes the natural aversion of mice for exposed and illuminated environments. The apparatus consists of a dark compartment (20 x 20 x 15 cm) with a stainless-steel grid floor and a small (5 x 5 cm) illuminated, elevated platform attached to its front center. During training, we placed each mouse on the platform and, as it stepped into the dark compartment, it received a foot-shock (0.3 mA, 5 Hz, 1 s). Forty-eight hours later, we placed each mouse on the platform again and the step-through latency was recorded (testing session, TS). If a mouse failed to cross within 300 s, the retention test was terminated and the mouse was assigned a ceiling score of 300. The mice were naïve to the dark compartment before the learning trial. When indicated, we conducted subsequent testing sessions at the designated time points for each experiment.

#### 4.4.2 Nose-poke habituation task

We also studied the nose-poke habituation task (HT) in a one-trial learning situation on a hole-board. The apparatus (Ugo Basile Mod. 6650, Comerio, Italy), made of a gray Perspex panel (40 x 40 x 22 cm), embodies 16 flush mounted tubes of 3 cm of diameter. Each tube has an infrared emitter and a diametrically opposed receiver connected to an automatic counter to register the number of nose-pokes into the holes. The training and testing trials were 24 h apart. During both trials, we placed each mouse at the center of the apparatus and the number of nose pokes was automatically registered for 5 min. From one mouse to the next, we carefully cleaned the hole-board with ethanol 70% to avoid olfactory cues.

### 4.5 Brain fixation and sectioning for Immunohistochemistry

Ninety minutes after the IA training, we deeply anesthetized animals with ketamine and xylazine, and fixed their tissue by intracardiac perfusion of ~10 mL of ice-cold 0.1% heparin phosphate buffered saline (PBS), followed by ~25 mL of ice-cold 4% paraformaldehyde (PFA) in PBS. We then dissected their brains, which were post-fixed overnight in the same fixing solution. After 24 h, we first cryopreserved the tissue in a 15% sucrose solution in PBS at 4 °C for 24 h, then in a 30% sucrose solution in PBS for another 24 h and maintained it in a fresh 30% sucrose solution until sectioned. We embedded in Tissue-Tek (Sakura) and cryotome sectioned the brains at a thickness of 40 μm. Sections including CLA (−0.50 to 1.50 AP from bregma), CA1 area of the hippocampus (−2.50 AP to −1.50 from bregma) and motor cortices (0.00 to 1.50 AP from bregma; M1/2) were collected ^38^.

### 4.6 Immunohistochemistry (IHC)

We washed with 0.1% Triton in PBS and blocked the selected brain sections with 1% donkey serum (D9663, Sigma-Aldrich) in the wash solution for 60 min at room temperature (RT). Then, incubated the slices overnight at RT with 1:500 rabbit α c-Fos (ABE457, Merck Millipore) and 1:3000 mouse α parvalbumin (PV 235, Swant) antibodies. After 24 h, we washed them with 0.1% Triton in PBS, followed by incubation with 1:1000 Biotin donkey α mouse (715-065-151, Jackson ImmunoResearch). After 120 min, we again washed the sections with 0.1% Triton in PBS, then with PBS and lastly with phosphate buffer (PB), before incubating them with 1:200 donkey α rabbit Alexa 488 (711-545-152, Jackson ImmunoResearch) and 1:200 Streptavidin Cy3 (016-160-064, Jackson ImmunoResearch) in PB. After 120 min, brain sections were washed with PB, dried at RT, re-hydrated in PB and mounted in gelatinized slides (gelatine type a g8-500, Fisher Chemical) using Mowiol as mounting medium (81381, Sigma-Aldrich).

### 4.7 Acquisition and processing of images

We acquired images using a LSM900 Zeiss confocal microscope and a 20x objective (Carl Zeiss, Plan Apochromat, NA 0.8). Z-stacks spanning the entire thickness of each slice were obtained with 512 × 512 pixel resolution, parvalbumin (PV) and c-Fos acquisition channels were separated and images were processed with ImageJ. Briefly, we delineated a CLA-mask using the signal present in the PV channel. Then, the signal from c-Fos channel was collapsed to a maximum intensity projection and the CLA-mask superimposed. We normalized the amount of c-Fos positive cells to the mask area for each image. CA1 and M1/2 masks were performed based on the atlas of Franklin and Paxinos ^38^.

### 4.8 Data analyses

We show data representing latency to step-through, cumulative nose pokes and relative density of c-Fos positive cells in plots to depict summary statistics, as well as the distribution of individual values. For normally distributed data, we depict the mean and standard error as summary statistics, otherwise we show the median and interquartile range (IQR). The summary statistics and final amount of mice included in the analysis of each experiment is specified in its respective Figure legend. We performed data analysis with R (version 4.2.2). To account for data dependency (as multiple observations were drawn from each animal), generalized linear mixed models (GLMMs) were performed using the *glmmTMB* package (version 1.1.6)^42^. For behavioral data, we fitted a GLMM for each variable of interest with Group (Veh and Lido) and Trial (TR, TS1, TS2 and TS3 in Figures) as fixed effects, and mouse ID (Subject) for random effects. We added the Minute fixed effect (1 through 5 or 15) for HT experiments. For IHC data, we fitted a GLMM for c-Fos density in each image of CLA, CA1 or M1/2 with Group (Na and TR) as fixed effect and mouse ID (Subject) for random effect (ANOVA of GLMMs are displayed in Suppl. Table 1). The GLMM corresponding to the latency to step-through (IA experiments) was fitted with a Gamma family distribution and a log link function, while the LMM for cumulative nose pokes (HT experiments) and IHC experiments were fitted with a Gaussian family distribution and an identity link function. The goodness-of-fit for each model was analyzed using the *DHARMa* package (version 0.4.6). Visual inspection of residual plots did not reveal any concerning deviations from normality or homoscedasticity. When applicable, we performed post-hoc comparisons using the *emmeans* package (version 1.8.5)^43^ and adjusted all reported p-values for multiple comparisons with the Šidák correction method. In all cases, p values under 0.05 were considered significant. When indicated, 1 signifies p < 0.05; 11, p < 0.01; and 111, p < 0.001.

## Author Contributions

MMB, CM, SOR and MCK contributed to the study conception and design, material preparation, data collection and analysis. Visualization of data was performed by SOR and CM. MMB supervised the project and secured the required funding for the project. The development of methodology and provision of resources were performed by AGR, MMB and AMD. The first draft of the manuscript was written by MMB and CM, and all authors commented on previous versions of the manuscript. All authors read and approved the final manuscript.

## Data Availability

The datasets generated during and analyzed during the current study are available in a permanent repository.

## Ethics approval

This study was carried out in accordance with the National Institute of Health Guide for the Care and Use of Laboratory Animals (NIH Publication N1 80–23/96) and approval was granted by the local research ethics committee (CICUAL-FFyB, CUDAP # 0044975-2016). All efforts were made to minimize animal suffering and to reduce the number of animals used. This study is reported in accordance with ARRIVE guidelines.

## Abbreviations

CLA: claustrum
GLMM: generalized linear mixed model
HT: nose-poke habituation task
IA: inhibitory avoidance
IHC: immunohistochemistry
IQR: interquartile range
Lido: lidocaine
M1/2: motor cortices
Na: Naïve
PB: phosphate buffer
PBS: phosphate buffered saline
PFA: paraformaldehyde
PV: parvalbumin
RT: room temperature
TR: training
TS: testing
Veh: vehicle

## Acknowledgments

MCK and MMB are members of CONICET and CM is a postdoctoral fellow of CONICET. We would like to thank Dr. Joaquín Piriz for generously providing us with fluorescent beads, which were instrumental in the setup of the intraclaustral surgery. We also thank Dr. Juan Belforte for kindly providing us with the mouse α parvalbumin antibody, an invaluable tool that aided in the morphological delineation of the CLA.

