## Supplementary material for "Delving into the claustrum: insights into memory formation, stabilization and updating in mice": Suppl. Fig. 1 and Table 1

### Supplementary data

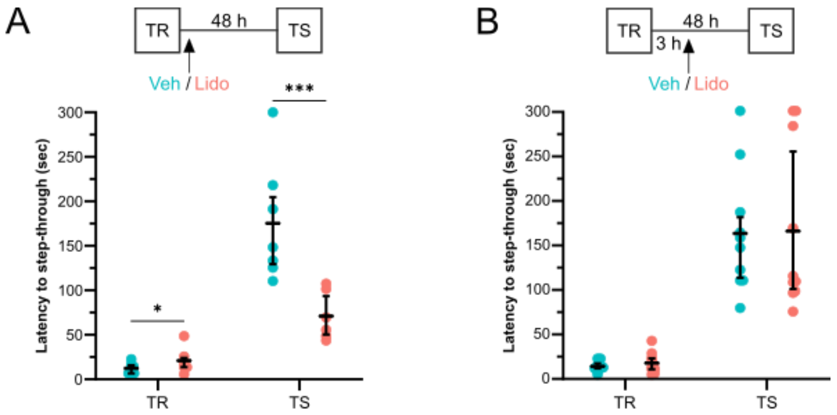


**Supplementary Figure 1.** The claustrum in the acquisition and consolidation of an associative memory in female mice. **(A-B, top panel)** We injected mice in the CLA with Veh or Lido solution either **(A)** immediately after TR ($n_{Veh}$ = 7, $n_{Lido}$ = 6) or **(B)** 3 h after TR ($n_{Veh}$ = 10, $n_{Lido}$ = 10). The retention test took place 48 h later (TS). **(A-B, bottom panel)** Step-through latencies of animals during TR and TS trials. Median values and IQR are shown in black, individual values in color. Veh vs Lido: **(A)** p = .0352 during TR, p = .0005 during TS; **(B)** p = .2722 during TR, p = .9470 during TS.

| Figure number | Experiment number and n | Fixed factor | df | $\chi^2$ | p value | Sign. |
|---|---|---|---|---|---|---|
| 1 | 1 - CLA<br>$n_{Na}$ = 6-13 (4)<br>$n_{TR}$ = 2-14 (4) | G | 1 | 8.20 | .0042 | ** |
| | 1 - CA1<br>$n_{Na}$ = 2-6 (4)<br>$n_{TR}$ = 3-6 (4) | G | 1 | 13.51 | .0002 | *** |
| | 1 - M1/2<br>$n_{Na}$ = 7-12 (4)<br>$n_{TR}$ = 7-14 (4) | G | 1 | 1.47 | .2249 | ns |
| 2 | 2<br>$n_{Veh}$ = 9<br>$n_{Lido}$ = 8 | G | 1 | 33.72 | 6.349e-09 | *** |
| | | T | 1 | 231.30 | < 2.2e-16 | *** |
| | | G*T | 1 | 5.92 | .0149 | * |
| | 3<br>$n_{Veh}$ = 8<br>$n_{Lido}$ = 7 | G | 1 | .16 | .6908 | ns |
| | | T | 1 | 1385.09 | <2e-16 | *** |

| | | | | | | |
|---|---|---|---|---|---|---|
| | | G*T | 1 | .0026 | .9593 | ns |
| | 4<br><br>$n_{Veh}$ = 8<br>$n_{Lido}$ = 9 | G | 1 | .03 | .8622 | ns |
| | | T | 1 | 243.26 | <2e-16 | *** |
| | | G*T | 1 | .11 | .7420 | ns |
| 3 | 5<br><br>$n_{Veh}$ = 6<br>$n_{Lido}$ = 5 | G | 1 | .71 | .3988 | ns |
| | | T | 1 | 123.84 | < 2.2e-16 | *** |
| | | M | 1 | 857.79 | < 2.2e-16 | *** |
| | | G*T | 1 | 28.65 | 8.65e-08 | *** |
| | | T*M | 4 | 30.69 | 3.54e-06 | *** |
| | | G*T*M | 4 | 14.88 | 0.0049 | ** |
| | 6<br><br>$n_{Veh}$ = 7<br>$n_{Lido}$ = 11 | G | 1 | .84 | .3580 | ns |
| | | T | 1 | 88.18 | < 2.2e-16 | *** |
| | | M | 1 | 560.30 | < 2.2e-16 | *** |
| | | G*T | 1 | 1.40 | .2367 | ns |
| | | T*M | 4 | 20.37 | .0004 | *** |
| | | G*T*M | 4 | 2.88 | .5788 | ns |
| | 7<br><br>$n_{Na}$ = 7<br>$n_{Veh}$ = 7<br>$n_{Lido}$ = 6 | G | 2 | 1.34 | .5113 | ns |
| | | T | 14 | 3179.71 | <2e-16 | *** |
| | | G*T | 28 | 23.98 | .6824 | ns |
| 4 | 8<br><br>$n_{Veh}$ = 13<br>$n_{Lido}$ = 12 | G | 2 | 1.80 | .1794 | ns |
| | | T | 14 | 641.66 | <2e-16 | *** |

| | | | | | | |
|---|---|---|---|---|---|---|
| | | G*T | 28 | 12.99 | .0047 | ** |
| | 9<br><br>$n_{Veh}$ = 8<br>$n_{Lido}$ = 6 | G | 1 | .002 | .9634 | ns |
| | | T | 1 | 251.27 | <2e-16 | *** |
| | | G*T | 1 | .32 | .5701 | ns |
| | 10<br><br>$n_{NR\text{-}Lido}$ = 9<br>$n_{TS1\text{-}Veh}$ = 9<br>$n_{TS1\text{-}Lido}$ = 10 | G | 1 | 2.02 | .3646 | ns |
| | | T | 1 | 459.71 | <2e-16 | *** |
| | | G*T | 1 | .93 | .6290 | ns |
| | 11<br><br>$n_{Veh}$ = 10<br>$n_{Lido}$ = 7 | G | 1 | .06 | .8009 | ns |
| | | T | 1 | 488.78 | <2e-16 | *** |
| | | G*T | 1 | 1.19 | .5522 | ns |
| Suppl. 1 | 12<br><br>$n_{Veh}$ = 7<br>$n_{Lido}$ = 6 | G | 1 | .93 | .334 | ns |
| | | T | 1 | 120.92 | <2e-16 | *** |
| | | G*T | 1 | 15.56 | 7.99e-05 | *** |
| | 13<br><br>$n_{Veh}$ = 10<br>$n_{Lido}$ = 10 | G | 1 | .68 | .41 | ns |
| | | T | 1 | 210.01 | <2e-16 | *** |
| | | G*T | 1 | .53 | .46 | ns |

**Table 1.** ANOVA results for GLMM models of each experimental dataset. The table shows the number of replicates corresponding to each experiment, as well as factor, degrees of freedom (df), Chi-square statistic ($\chi 2$), p value and significance of each ANOVA performed. G stands for Group factor, T for Trial and M for Minute. In experiment 1 (Figure 1), the number of replicates is noted as the amount of images of each mouse (the total number of mice in the experimental group).
